# Homozygote loss-of-function variants in the human *COCH* gene underlie hearing loss

**DOI:** 10.1101/2020.06.29.178053

**Authors:** Nada Danial-Farran, Elena Chervinsky, Prathamesh Thangaraj Nadar-Ponniah, Eran Cohen Barak, Shahar Taiber, Morad Khayat, Karen B. Avraham, Stavit A. Shalev

## Abstract

Since 1999, the *COCH* gene encoding cochlin, has been linked to the autosomal dominant non-syndromic hearing loss, DFNA9, with or without vestibular abnormalities. The hearing impairment associated with the variants affecting gene function has been attributed to a dominant-negative effect. Mutant cochlin was seen to accumulate intracellularly, with the formation of aggregates both inside and outside the cells, in contrast to the wild-type cochlin that is normally secreted. While an additional recessive variant in the *COCH* gene (DFNB110) has recently been reported, the mechanism of the loss-of-function (LOF) effect of the *COCH* gene product remains unknown. In this study, we used COS7 cell lines to investigate the consequences of a novel homozygous frameshift variant on RNA transcription, and on cochlin translation. Our results indicate a LOF effect of the variant and a major decrease in cochlin translation. This data has a dramatic impact on the accuracy of genetic counseling for both heterozygote and homozygote carriers of LOF variants in *COCH*.

## Introduction

The *COCH* gene encodes the secreted cochlin protein, a major component of the extracellular matrix of the inner ear [1]. Variants affecting *COCH* gene function have been linked to the autosomal dominant hearing loss, DFNA9 [2], and recently, also to the autosomal recessive hearing loss, DFNB110 [3, 4]. More than 20 dominant variants in the *COCH* gene are associated with progressive mild-to-profound hearing impairment and vestibular dysfunction in the second to seventh decade of life [5]. The majority of the variants are missense or in-frame variants. These variants are thought to be linked to dominant-negative or gain-of-function mechanisms that affect post-translation processing by disruption of the normal protein secretion, protein accumulation in the cells, or by generating protein dimers, leading to a damaging effect of the mutant protein [5]. A study of a frameshift variant in *COCH* further suggested that haploinsufficiency does not lead to hearing loss [6]. Despite this progress, loss-of-function (LOF) variants in the *COCH* gene have not been studied extensively.

In this study, we report the first homozygous frameshift variant in three hearing impaired family members of a Christian Arab family from northern Israel, identified using next-generation sequencing (NGS). We investigated RNA derived from the family, and post-translational events by expressing the wild-type and mutant constructs of *COCH* in cell lines, providing a functional analysis of LOF variants. In addition, we summarized the phenotypes in families with the recessive LOF variants in order to provide genotype-phenotype-correlations.

## Materials and methods

### Subjects

This study investigated a Christian Arab family from northern Israel with hearing impairment. All family members approved participation by signing informed consent forms, according to the instructions of the Ethical Committee of Ha’emek Medical Center, following the Helsinki Declaration. Detailed pedigrees, medical history, and audiograms were collected from deaf and hearing family members. Peripheral blood samples were obtained to extract genomic DNA and RNA.

### DNA sequencing

DNA extraction was performed using the FlexiGene DNA Kit (Qiagen, Germany). Sanger sequencing was used to rule out variants in the *GJB2* gene known to cause hearing loss and for variant segregation and screening (Table S1). Exome sequencing was performed for the deaf proband AF101-05. DNA sequencing was performed by Macrogen, USA. The library Agilent SureSelect v5 was employed, followed by NGS on the Illumina HiSeq4000, with an average read depth of 144X. The bioinformatics analysis was processed by the Variantyx Genomic Intelligence platform version 1.16.0.0

### RNA and reverse transcriptase-PCR

Total RNA extraction was performed using TRI reagent (Sigma). cDNA was prepared with 2 μg of total RNA using qScript^®^ (QuantaBio, USA). The mRNA expression of the wild-type and mutant alleles of *COCH* was examined using PCR amplification with tagged primers (Table S1).

### Plasmids and mutagenesis

For *COCH* expression, the human *COCH* cDNA sequence tagged with GFP (Sino Biological, Cat # HG11368ACG) was used. Mutagenesis was carried out by PCR to generate a mutant *COCH* cDNA.

### Cell culture

COS7 cells were cultured in Dulbecco’s Modified Eagle’s Medium (DMEM) with supplements, transfected using PolyJet™ (Signagen, Rockville, MD, USA), and fixed with 4% PFA. Dapi labelled the nuclei and m-Cherry-TNGP-N-10 (Addgene, Cat # 55145) localized the Golgi. Image acquisition was performed with a confocal microscopy system (LSM800 Carl Zeiss). The experiment was performed in triplicate.

### Western blotting

Cells were washed with PBS and lysed in urea sample buffer. Protein concentrations were equalized, samples were run on 7.5% SDS-PAGE gels. Gels were transferred to nitrocellulose membranes. Primary (anti-GFP, 1:2000, #A6455, Thermo Fisher Scientific, Waltham, MA, USA) and secondary antibodies (Goat anti-mouse and goat anti-rabbit peroxidase, 1:5000, #31430, #31460, Thermo Fisher Scientific) were used. Chemiluminescence was performed using the Syngene Imaging System (Yarden Biotech, Israel). Densitometry quantification was normalized using ⍰-actin (1:1000, #4970S, Cell Signaling Technologies, Danvers, MA, USA) as a loading control.

## Results

The family analyzed in this study has three affected children (Fig. 1a). The hearing impairment is similar in all affected family members who have high-tone hearing loss, ranging from mild-moderate in low frequencies to severe in high frequencies (Fig. 1b). The proband, 101-05, was diagnosed at the age of two years, as her parents noticed delayed lingual development. Diagnosis of the hearing disability was confirmed with auditory brainstem response (ABR) tests. Family members 101-06 and 101-07 were diagnosed prior to one year of age. As the hearing loss of proband 101-05 is mild-to-severe, and was diagnosed by the age of 2, it may have been congenital but diagnosed later due to lack of awareness. This assumption is supported by the fact that after the diagnosis of the first affected child, the two other affected children were diagnosed at the age of a few months.

**Fig. 1.**
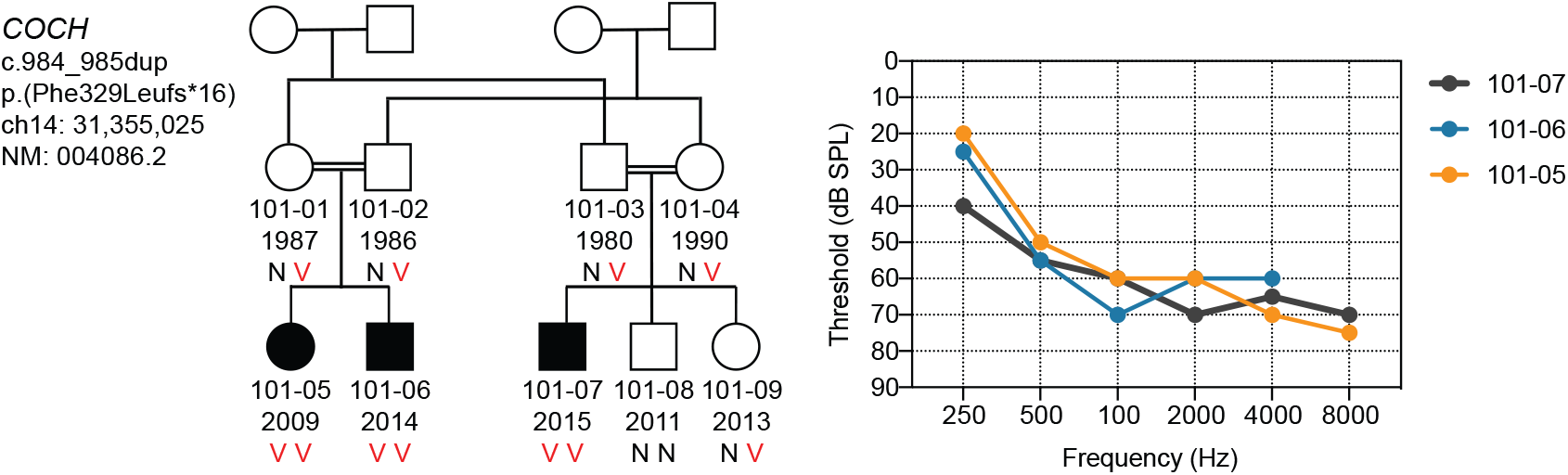
Family AF101 pedigree, segregation of the *COCH* variant and audiograms. (a) The AF101 family pedigree, showing multiple family members with congenital, or pre-lingual onset, mild-moderate down-sloping to severe high-tone hearing loss. N, normal allele; V, variant allele. (b) Audiograms of individuals 101-05, 101-06, and 101-07 indicate moderate to severe hearing impairment.

WES was performed on proband 101-05 and generated 109 million reads, with an average coverage of 144X. The sequencing analysis revealed over 150K variants. Our interpretation of the variants focused on homozygous alleles and is based on the assumption of shared ancestral pathogenic alleles due to consanguinity in the family. Variants in databases, including gnomAD with MAF > 0.01, were excluded. The remaining variants were investigated for their potential damaging effect on the auditory system. The variant c.984_985dup in the *COCH* gene, which does not appear in the gnomAD database, was selected as the lead candidate. Of the 54 LOF variants in the *COCH* gene reported in the gnomAD database, none were in the homozygous state. In addition, there were fewer LOF variants than the expected LOF variant number, according to the gnomAD depth corrected probability of this type of variant for each gene [7]. The WES analysis was followed by Sanger sequencing (Table S1), which confirmed co-segregation of the variant with hearing impairment in an autosomal recessive manner, for the nine family members examined (Fig. 1a). No carriers were detected among 110 hearing Christian Arab adults living outside the family’s village; five carriers were detected among 100 hearing Christian Arab controls living in the same village.

To study the expression of the *COCH* mutant allele, RNA was extracted from two heterozygous parents and three homozygous children and cDNA was prepared by reverse transcriptase-PCR (RT-PCR). Gel electrophoresis of the PCR products indicated no differences in the expression of the wild-type and mutant *COCH* alleles (Fig. S1a, b). The expression of both alleles was validated by fragment-analysis PCR with fluorescent primers (Fig. S1c, d). This is in contrast to the previously reported variant, c.292C>T, p.Arg98*, which was linked to activating nonsense mediated decay (NMD), where the mutant allele was not detectable [3]

To investigate the consequences of protein expression, and possible accumulation of the mutant protein in the cells, we co-transfected the COS7 cell line with COCH-GFP plasmids containing both the wild-type and mutant forms of the cDNA, and a vector expressing TGNP-mCherry, a Golgi complex protein. Wild-type cochlin was detected in the cellular cytoplasm and co-localized in the Golgi complex, as previously reported [8]. GFP fluorescence could not be detected in cells transfected with the mutant COCH (Fig. 2a). This finding was also confirmed by western blot, in which we used an anti-GFP antibody to detect cochlin (Fig. 2b). Protein lysate from cells transfected with wild-type *COCH* showed a clear band at 95kD, as expected, while mutant *COCH* could not be detected.

**Fig. 2.**
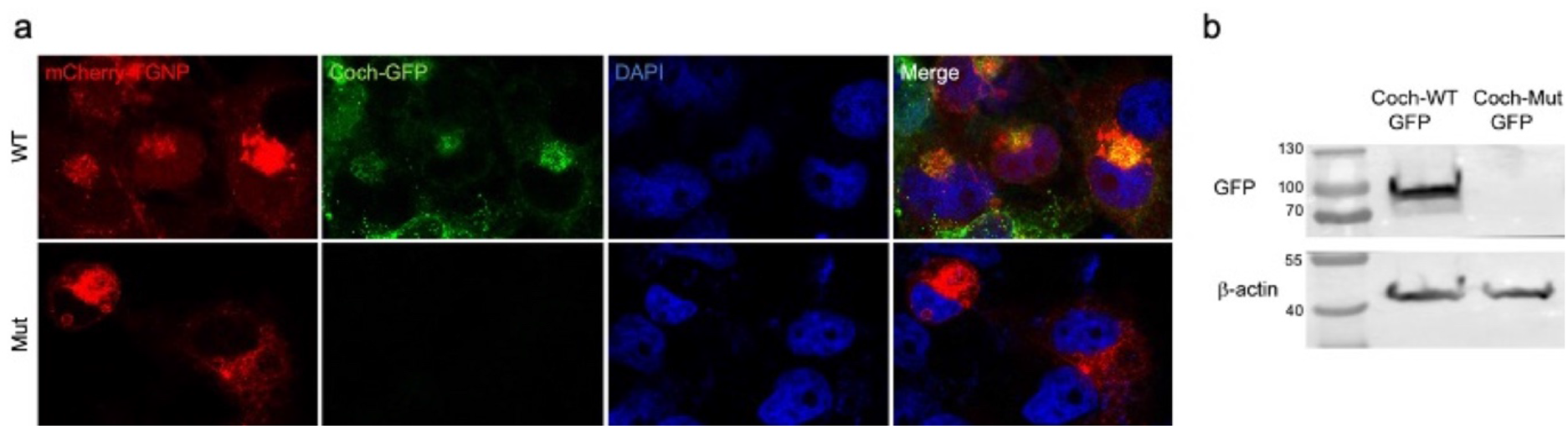
The c.984_985dup variant results in loss of cochlin protein expression. (a) Imaging of COS7 cells transfected with wild-type and mutant forms of COCH-GFP. Mutant cochlin was not detected in the cells transfected with the variant. (b) Cochlin protein expression was only detected in the cells transfected with wild-type COCH-GFP, while the β-actin, serving as a control, was observed in both cells.

## Discussion

Exome sequencing allowed us to identify the first homozygote frameshift variant in the *COCH* gene, which segregates with hearing loss in a Christian Arab family living in northern Israel. This variant is the third LOF allele reported to cause early onset sensorineural hearing loss in a homozygote state in the *COCH* gene. The three families (Table 1) are from different origins, all reported endogamous marriages, with a total of seven affected individuals. Notably, the reported patients exhibit high-tone down-sloping from mild-moderate to severe hearing loss. In the dominant pathogenic alleles, damage is first seen at the higher frequencies and then affects mid and low frequencies [2, 9, 10]. In contrast, the pathogenic dominant alleles tend to lead to late-onset hearing impairment [5]. The dominant pathogenic alleles of *COCH* gene mainly disrupt the post-translational processes in the cell and interrupt secretion to the extracellular matrix [5]. Conversely, the homozygous *COCH* alleles affect hearing by a dramatic reduction in the amount of the cochlin protein product, in accordance with a LOF mechanism.

**Table 1.** Clinical data of hearing-impaired individuals with homozygote LOF variants.

| *Family* | *Ethnicity* | *Affected individual* | *Onset* | *Pure tone audiometry (PTA; 0.5, 1, 2kHz), Age* | *Progression of hearing loss severity\** | *TEAOE* | *DPOAE* | *Vestibular dysfunction* | *Genomic coordinate (GRCH37)* | *cDNA position (RefSeq NM: 004086.2; NG_008211.2)* | *Predicted effect on protein* | *Molecular analysis* | *References* |
| --- | --- | --- | --- | --- | --- | --- | --- | --- | --- | --- | --- | --- | --- |
| AF101 | Christian Arab | 101-05 | Congenital | R 60 dB, 7<br>L 63 dB, 7 | Yes | Absent | Absent | No | g.31,355,025 | c.984_985dup | p.(Phe329fs*16) | No NMD activation<br>Reduced protein translation | This study |
|  |  | 101-06 | Congenital | R 51 dB, 4<br>L 62 dB, 4 | Yes | NA | NA | No |  |  |  |  |  |
|  |  | 101-07 | Congenital | R 53 dB, 4<br>L 62 dB, 4 | NA | NA | NA | No |  |  |  |  |  |
| AI | Moroccann | AI-010 | Congenital | R 63 dB, 4<br>L 68 dB, 4 | Yes | Absent | Absent | Yes | g.31,348,069 | c.292C>T | p.(Arg98Ter) | NMD activation | [3] |
|  |  | AI-011 | Congenital | R 55 dB, 0.7<br>L 57 dB-0.7 | NA | Absent | Absent | NA |  |  |  |  |  |
| 9600236 | Iranian | IV-1 | Prelingual | R 67 dB, 35<br>L 70 dB, 35 | NA | NA | NA | No | g.31,346,811 | c.116T>A | p.(Leu39Ter) | NA | [4] |
|  |  | IV-6 | Prelingual | NA | NA | NA | NA | No |  |  |  |  |  |
|  |  | IV-7 | Prelingual | NA | NA | NA | NA | No |  |  |  |  |  |

NA, not available. L, left. R, right

The nonsense variants reported in the Iranian and Moroccan families are located in the second and third exon of the *COCH* gene (NG_008211.2), respectively, which may explain the activation of NMD as a result of the nonsense variant c.292C>T in the Moroccan family [3], unlike the variant detected in this study, located at exon 9 out of 11 exons (NG_008211.2). Investigating the *in vivo* expression of the wild-type and mutant alleles of the *COCH* gene, by examining mRNA extracted from leukocytes, excluded the option of NMD activation. Our results indicate an equal level of mRNA expression of the wild-type and mutant allele. Next, we compared the translation of the mutant to that of the wild-type allele in COS7 cells transfected with plasmids expressing both wild-type and mutant forms of the cochlin protein. The data revealed a dramatic reduction of the mutant cochlin, compared to the wild-type, which was clearly detected by green fluorescence in the GFP-transfected cells (Fig. 2a), and confirmed by western blot (Fig. 2b). These results support a causal effect of LOF alleles that reduce protein translation, lead to reduced stability, or facilitate degradation [11].

In addition to the immediate impact of these findings on clinical genetics, the data from our study provide a better understanding of cochlin function and validate the LOF mechanism of the *COCH* recessive variants.

## Data availability

The c.984_985dup variant is available on ClinVar (Accession # VCV000870100).

## Acknowledgements

We thank the families for their participation in this study. We also thank Dr. Zippora Brownstein for helpful comments throughout the research.

## Supplementary information

**Fig. S1.**
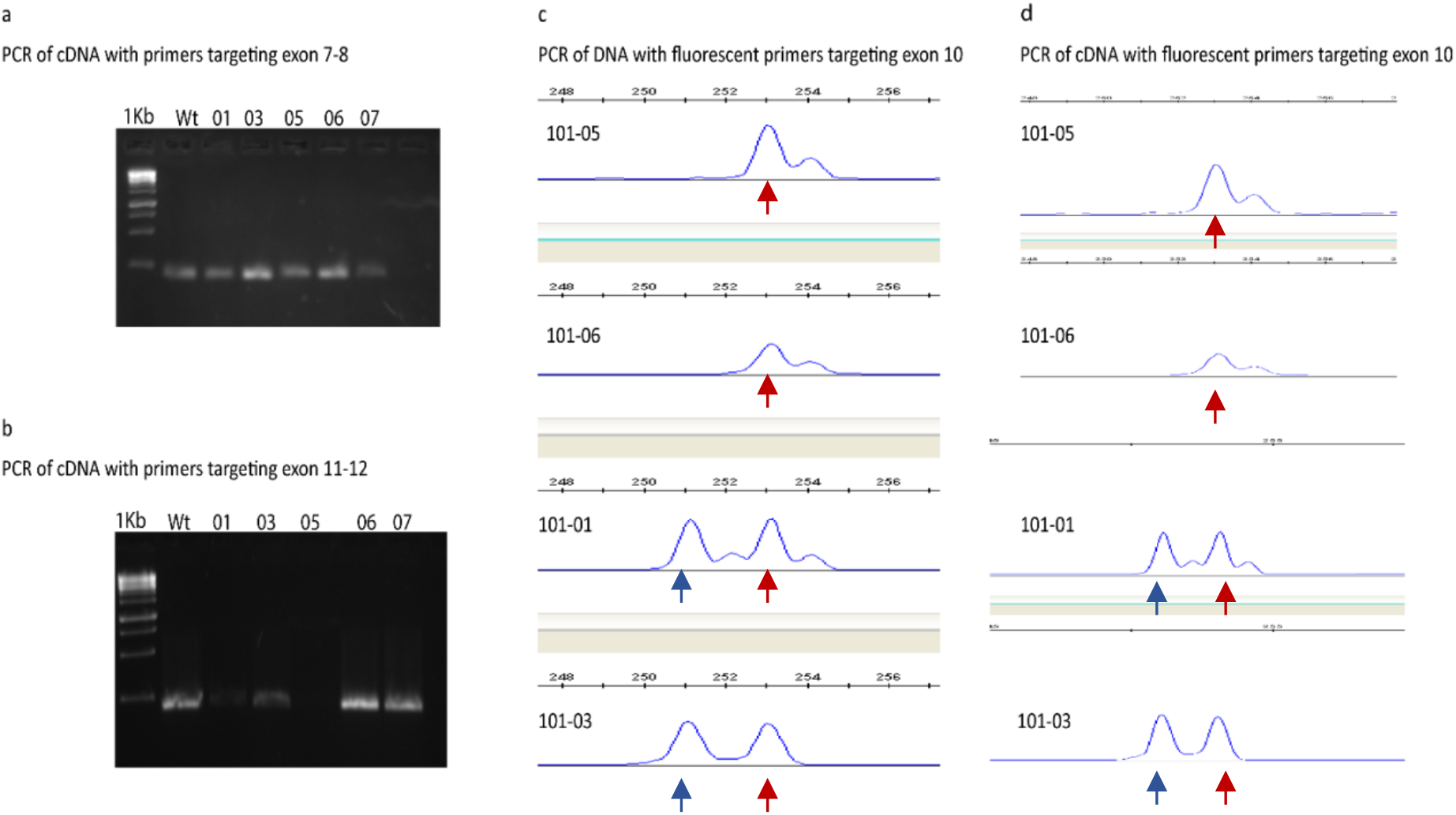
Expression analysis of the COCH wild-type and mutant alleles. **a, b** Agarose gel electrophoresis of PCR products of cDNA synthesized from RNA extracted from family members with primers targeting an upstream variant sequence (a), and downstream of the variant (b). **c** Fragment analysis of DNA for parents and two affected siblings. **d** Fragment analysis of cDNA for parents and two affected siblings. The blue arrow indicates the wild-type allele and the red arrow indicates the mutant allele.

**Table S1.**
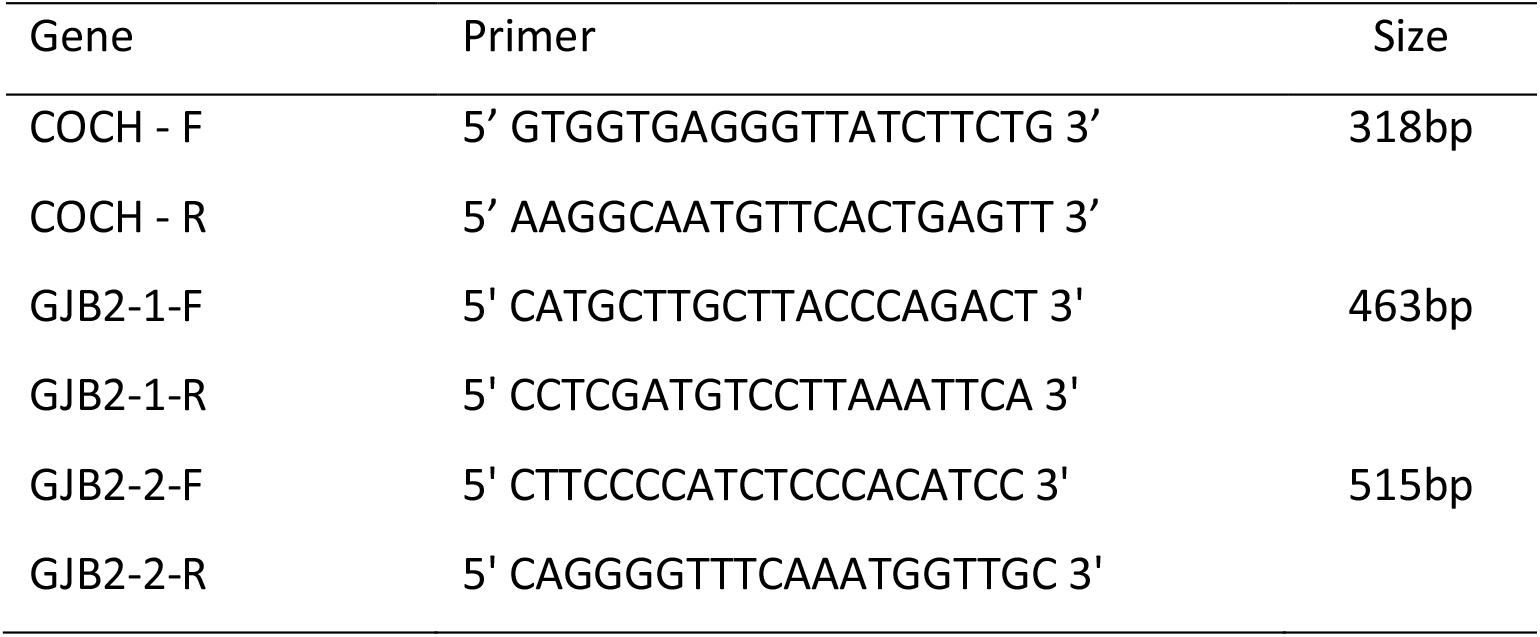
Primers used for screening of genes.

